# Competitive advantage of hypervirulent Group B *Streptococcus* for neonatal colonization revealed by metagenomic analysis of gut microbiota

**DOI:** 10.64898/2026.06.18.733191

**Authors:** Anne-Sophie Alvarez, Florian Plaza Oñate, Gérald Touak, Sean Kennedy, François Goffinet, Céline Plainvert, Laurent Mandelbrot, Stanislav Dusko Ehrlich, Claire Poyart, Asmaa Tazi

## Abstract

Group B *Streptococcus* (GBS) is the leading cause of neonatal invasive infections. Late-onset infections (7-89 days after birth) are caused by GBS clonal complex 17 (CC17) in 50-80% of cases, likely resulting from bacterial translocation across the intestinal barrier. However, the factors influencing GBS colonization in neonates are incompletely understood. We used shotgun metagenomics on fecal samples from 100 neonates aged 21 days and identified taxonomic signatures of GBS colonization, including decreased *Enterobacter hormaechei* abundance in neonates colonized by non-CC17 GBS. Using *in vitro* assays with representative isolates, we demonstrate that GBS CC17 competes more effectively than GBS non-CC17 against *E. hormaechei,* with enhanced adherence to enterocytes mediated by the CC17-specific HvgA adhesin. Our findings highlight lineage-dependent interspecies interactions of GBS that likely influence its ability to colonize the neonatal gut. These interactions must be considered when developing microbiota-based strategies to mitigate neonatal colonization and infection by GBS.

## Introduction

*Streptococcus agalactiae* (Group B *Streptococcus*, GBS) is a Gram-positive encapsulated bacterium that commonly colonizes the intestinal and vaginal tracts of 10 to 30% of healthy adults.^1^ However, despite its frequent asymptomatic carriage, GBS is also the leading cause of bacterial invasive infections in neonates worldwide.^2^ Neonatal GBS infections primarily manifest as sepsis and, in 10–30% of cases, as meningitis, and remain associated with a substantial burden, with an estimated mortality rate of 10% and long-term sequelae in approximately 30% of cases.^3–5^

Neonatal GBS infections are divided into early-onset disease (EOD) and late-onset disease (LOD), which occur during the first week of life and between 7 and 89 days of life, respectively.^6^ These two syndromes differ not only in chronology but also in pathophysiology. In EOD, neonates become infected by inhalation or ingestion of contaminated amniotic fluid or maternal vaginal secretions prior to or during parturition. Conversely, the mechanisms underlying neonatal colonization and infection in LOD remain incompletely understood, although the gastrointestinal tract represents the most likely portal of entry.^7–9^

Worldwide molecular epidemiological studies demonstrated that a substantial proportion of EOD cases and the vast majority of LOD cases are attributable to a specific GBS clonal complex (CC) almost exclusively of capsular type III and designated the hypervirulent GBS CC17.^10–13^ The hypervirulence of GBS CC17 has been attributed to the expression of two specific surface proteins, HvgA and Srr2, that increase gut colonization in mouse models as well as GBS translocation across the intestinal and blood-brain barriers.^8,9^ Furthermore, GBS CC17 exhibits a greater capacity to colonize the infant gut compared to other GBS clones, as demonstrated in the human Col-StreptoB cohort,^14^ suggesting that this property could contribute to its propensity to cause LOD. Nevertheless, the factors influencing GBS establishment in the gut during the early weeks of life, beyond maternal colonization, remain poorly understood. Among these, neonatal gut microbiota and diet could play an important role.^15^

In this study, we investigated the relationship between neonatal gut microbiota and GBS colonization in 100 neonates aged 21 ± 7 days, i.e. at an age coinciding with the median age of LOD onset.^2^ We integrated fecal metagenomics, clinical variables, including mode of delivery and feeding diet, and *in vitro* models of interspecies bacterial competition to decipher GBS-microbiota interactions involved in neonatal gut colonization by GBS. Our findings highlight lineage-specific signatures of GBS colonization and demonstrate a competitive advantage of GBS CC17 compared to other lineages that is associated with the CC17-specific HvgA adhesin. Together, these results provide a mechanistic framework for GBS colonization dynamics in the neonatal gut and support the development of microbiota-informed strategies to reduce neonatal GBS colonization and infection.

## Results

### Study design and cohort characteristics

To identify the fecal microbiota signatures associated with neonatal GBS colonization, we analyzed a subset of 100 neonates aged 21 ± 7 days old from the prospective Col-StreptoB cohort study conducted in France between 2012 and 2015 (ClinicalTrials.gov #NCT01719510). The study aimed to identify demographic, clinical, and risk factors of GBS colonization in neonates and infants (see the Methods section). Eligible participants were pregnant women over 18 years old who were screened positive for vaginal GBS colonization. Mother-infant pairs were recruited and monitored after maternity discharge at 21 ± 7 days and 60 ± 7 days. A total of 890 mother-infant pairs were included, and 748 of them completed follow-up at 21 ± 7 days. Among those, 157 neonates (21%) were found colonized by GBS, as determined by conventional culture methods of pharyngeal and fecal samples.^14^ None of the infants were diagnosed with a GBS disease.

The analyzed subset comprised three groups: GBS non-colonized, GBS non-CC17-colonized, and GBS CC17-colonized neonates, as defined by culture methods and molecular typing. The subset included all neonates positive for GBS CC17 (n=10), a random selection of neonates positive for GBS non-CC17 (n=40), and a random selection of neonates negative for GBS (n=52) (Figure 1). Shotgun metagenomic sequencing was performed on fecal samples, and two samples from non-CC17-colonized neonates were excluded due to poor sequencing quality, resulting in a total of 100 analyzed samples. The baseline and follow-up characteristics of the studied population are summarized in Table 1. Notably, vaginal delivery was largely predominant (82%) and 95% of mothers received intrapartum antibiotic prophylaxis (IAP), as recommended by the French guidelines for the prevention of GBS EOD.^16,17^ Key potential confounding variables for microbiota composition, including mode of delivery, gestational age at delivery, birthweight, antibiotic intake prior to delivery and until 21 ± 7 days of age, duration of IAP, and feeding diet, did not differ significantly among the three groups of neonates (Table 1).

**Figure 1.**
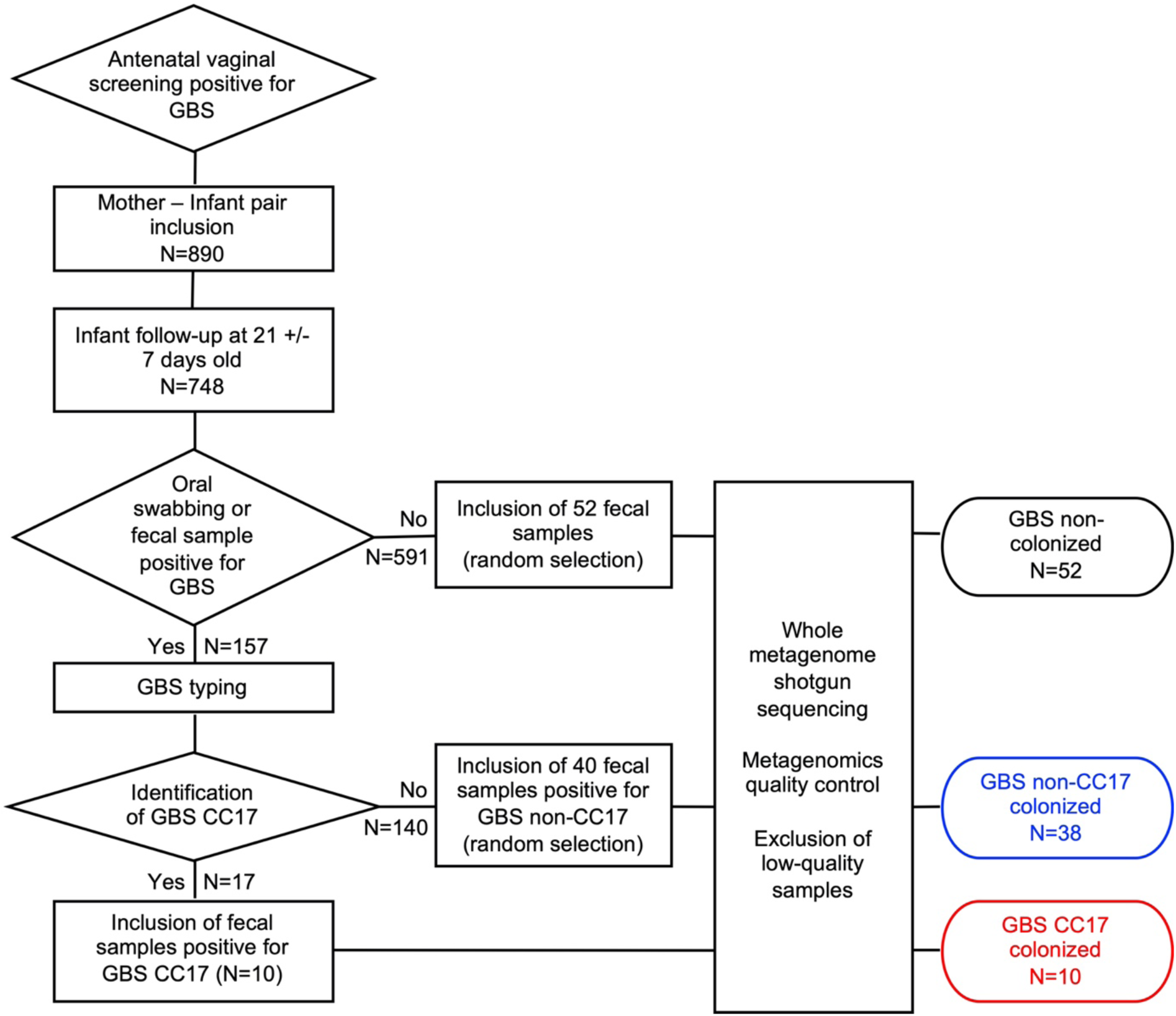
Study flow diagram and inclusion criteria of stool samples for metagenomic analysis. CC17: clonal complex 17; GBS: Group B *Streptococcus*.

**Table 1.** Descriptive statistics of the study population. ^a^ including 2 patients from Mauritius, 3 from Asia, 1 from Sri-Lanka, and 1 from Brazil. CC: clonal complex; GBS: Group B *Streptococcus;* NICU: neonatal intensive care unit; ROM: rupture of membranes. SD: standard deviation. Statistical analysis was performed using Chi-square or Fisher exact tests for categorical variables and one-way-ANOVA for continuous variables.

| Colonization by GBS at 21 +/-7 days old | No GBS<br>N=52 | GBS non-CC17<br>N=38 | GBS CC17<br>N=10 | p-value |
| --- | --- | --- | --- | --- |
| Neonatal sex ratio M/F | 1.2 | 1.4 | 1.0 | 0.90 |
| Maternal age, years (mean +/- SD) | 32.2 +/- 4.4 | 31.4 +/- 4.3 | 32.6 +/- 3.0 | 0.60 |
| Maternal place of birth |  |  |  | 0.86 |
| Europe | 27 (54.0%) | 23 (62.2%) | 5 (55.6%) |  |
| North Africa | 14 (28.0%) | 6 (16.2%) | 2 (22.2%) |  |
| Sub-Saharan Africa | 5 (10.0%) | 6 (16.2%) | 1 (11.1%) |  |
| Others<sup>a</sup> | 4 (8.0%) | 2 (5.4%) | 1 (11.1%) |  |
| Missing | 2 (3.8%) | 1 (2.6%) | 1 (10.0%) |  |
| Pregnancy associated complications |  |  |  | 0.99 |
| Hospitalization for threatened preterm labor | 0 | 0 | 0 |  |
| Gestational diabetes | 7 (13.5%) | 4 (10.5%) | 1 (10%) |  |
| Hypertension | 2 (3.8%) | 1 (2.6%) | 0 |  |
| ROM before entry in delivery room | 16 (30.8%) | 16 (42.1%) | 4 (40%) | 0.52 |
| Mode of delivery |  |  |  | 0.77 |
| Vaginal | 41 (78.8%) | 33 (86.8%) | 8 (80%) |  |
| Caesarean in labor | 8 (15.4%) | 4 (10.5%) | 2 (20%) |  |
| Planned caesarean | 3 (5.8%) | 1 (2.6%) | 0 |  |
| Gestational age at delivery, weeks (mean +/- SD) | 39.6 +/- 1.1 | 39.6 +/- 1.2 | 39.8 +/- 1.1 | 0.87 |
| Birthweight, grams (mean +/- SD) | 3451 +/- 445 | 3514 +/- 412 | 3408 +/- 412 | 0.70 |
| Transfer to NICU, n (%) | 2 (3.8%) | 1 (2.6%) | 2 (20%) | 0.07 |
| Antibiotics in the month before delivery | 0 | 0 | 0 | - |
| Antibiotics before 21 +/- 7 days of life | 5 (9.6%) | 1 (2.6%) | 0 | 0.27 |
| Intrapartum antibiotic prophylaxis |  |  |  |  |
| Duration, hours, median (range) | 5.02 (0 – 43.78) | 6.23 (0 – 13.22) | 5.11 (0 – 8.45) | 0.34 |
| None | 4 (7.6%) | 1 (2.6%) | 0 | 0.57 |
| < 4h | 16 (30.8%) | 10 (26.3%) | 4 (40%) |  |
| ≥ 4h and < 8h | 16 (30.8%) | 16 (42.2%) | 5 (50%) |  |
| ≥ 8h | 16 (30.8%) | 11 (28.9%) | 1 (10%) |  |
| Weight at 21 +/- 7 days, grams (mean +/- SD) | 3948 +/- 559 | 3966 +/- 526 | 3919 +/- 548 | 0.97 |
| Feeding diet at 21 +/- 7 days of life |  |  |  | 0.15 |
| Breastfeeding | 26 (52.0%) | 19 (52.8%) | 6 (60%) |  |
| Formula feeding | 16 (32.0%) | 5 (13.9%) | 1 (10%) |  |
| Mixed diet | 8 (16.0%) | 12 (33.3%) | 3 (30%) |  |
| Missing | 2 (3.8%) | 2 (5.3%) | 0 |  |

### Neonatal GBS colonization is not associated with fecal microbiota α diversity and overall composition

To address potential associations between GBS intestinal colonization and microbiota composition, sequencing reads from each fecal sample were first aligned against a gene catalog representative of the neonatal gut microbiota.^18,19^ Mapping rates were high (79.1% ± 3.6%), close to saturation, and similar between the three groups (Figure 2a). MetaGenomic Species (MGS) richness and Shannon diversity index did not differ between groups (Figures 2b and 2c), suggesting no association between GBS colonization and gut microbiota α diversity at 21 ± 7 days of life.

**Figure 2.**
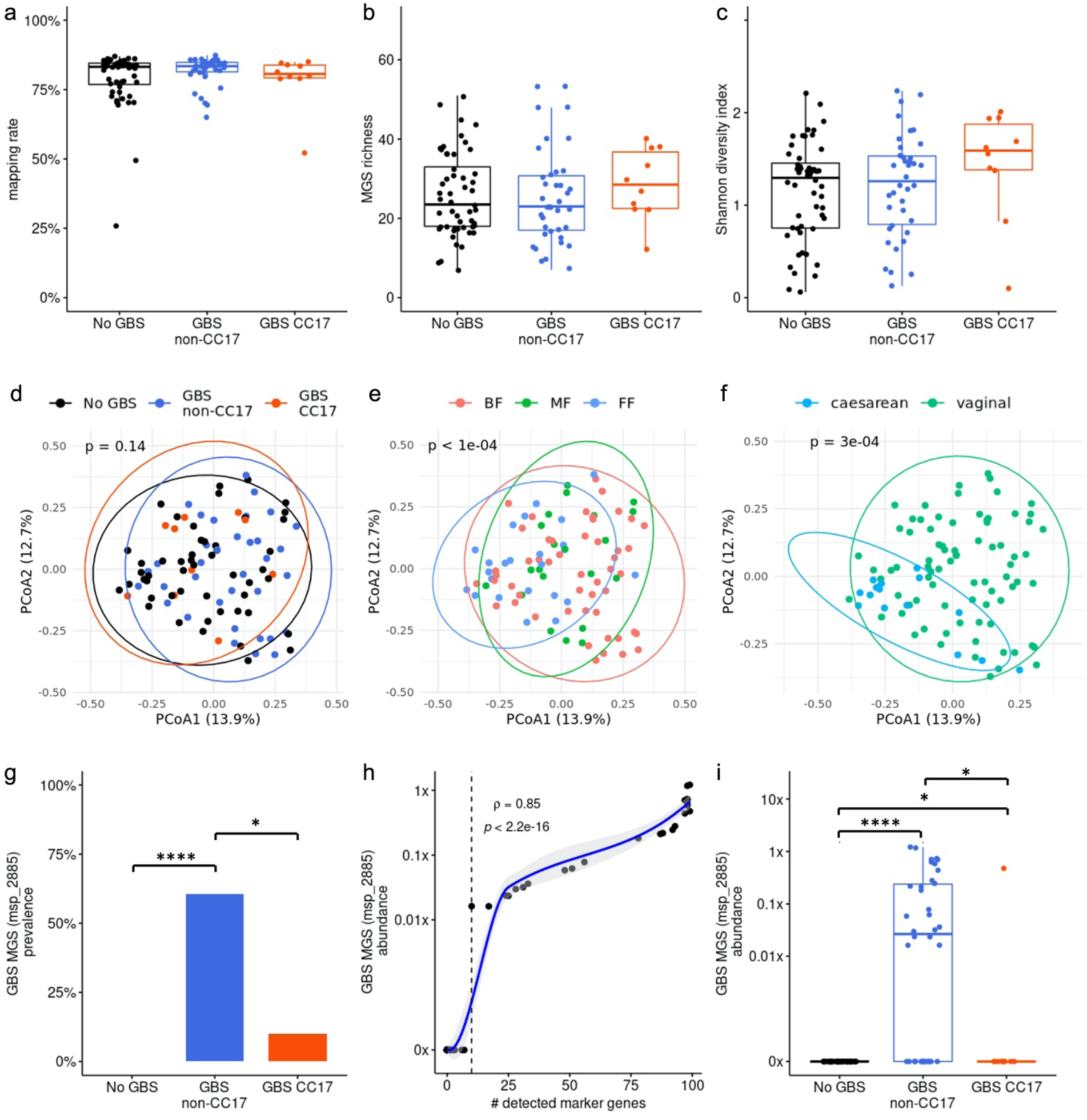
General features of fecal microbiota composition according to GBS colonization in 21 ± 7-day-old neonates. (a-c) Mapping rates (a), MGS richness (b) and Shannon diversity index (c). (d-f) Principal coordinates analysis (PCoA) based on Bray-Curtis dissimilarity at MGS level showing group differences in community structure according to GBS colonization status (d), neonatal diet (e), and mode of delivery (f). Prevalence (g) and abundance (h-i) of the MGS msp_2885 representative of the GBS species according to the number of detected marker genes (h) and to GBS colonization status (i). (a-c, i) Data are displayed as box-and-whisker plots representing the interquartile range, with the central line indicating the median; individual data points are shown as dots. Statistical analyses were performed using the Kruskal Wallis test followed by the Wilcoxon rank sum test (a-c, i), Permutational Multivariate Analysis of Variance (PERMANOVA) (d-f), Fisher’s exact test (g), and Spearman’s correlation (h). * p<0.05; **** p<0.0001. BF: breastfeeding; CC: clonal complex; FF: formula feeding; GBS: Group B *Streptococcus*; MF: mixed feeding; MGS: metagenomic species.

Next, we investigated the impact of GBS colonization on overall microbiota composition using permutational multivariate analysis of variance (PERMANOVA) based on Bray–Curtis dissimilarity at the MGS level. We found no significant association between fecal microbiota composition and GBS colonization status (p=0.14, Figure 2d). In contrast, the same analysis confirmed the importance of diet, *i.e.* breastfeeding, formula feeding, and mixed feeding, and delivery mode in shaping gut microbiota composition in neonates (p<1.10^-4^ and p=3.10^-4^, respectively, Figures 2e and 2f).

### GBS abundance within the neonatal gut microbiota is low, especially in GBS CC17 carriers

Next, we assessed the prevalence and relative abundance of GBS within the neonatal gut microbiota. The MGS representative of GBS (msp_2885) was detected in only 50% (24/48) of the samples identified as positive for GBS by conventional culture methods and was absent from all culture-negative samples (Figure 2g). GBS-positive samples by culture but negative by metagenomics most likely contained GBS below the detection threshold, as an MGS was considered as present in a sample if at least 10 of its 100 marker genes were detected. Consistently, the number of detected marker genes was strongly correlated with the estimated GBS abundance (Spearman’s ρ = 0.85, Figure 2h). Based on the sequencing depth used in this study, we estimated that this detection threshold allowed reliable detection of GBS when its genome coverage exceeded 0.01x, which corresponds approximately to a relative abundance of 1/10 000.

Overall, the abundance of GBS was low (median genome coverage < 0.1x) as compared to that of *Escherichia coli* (median of approximately 10x), one of the earliest and most abundant colonizers of the neonatal gut (Figure 2i and Supplemental Figure 1). In addition, the abundance of the GBS MGS was higher in neonates colonized by non-CC17 strains compared to those colonized by CC17 strains (p=0.01, Figure 2i), indicating a heavier GBS intestinal colonization in non-CC17 carriers. Accordingly, the genes *srr1* and *bibA* that are harbored by non-CC17 GBS isolates were detected in 39% (15/38) of the samples from non-CC17-colonized neonates, whereas their GBS CC17 respective homologues *srr2* and *hvgA* were detected in only 10% (1/10) of the samples from CC17-colonized neonates.

### *Enterobacter hormaechei* abundance is reduced in neonates colonized by non-CC17 GBS

Metagenomic analysis identified a total of 147 bacterial genera across the 100 samples analyzed. Among these, 13 genera belonging to four major phyla (*Actinomycetota*, *Bacteroidota*, *Bacillota*, and *Pseudomonadota*) were predominant with a mean relative abundance exceeding 1% (Figure 3a). Six genera differed significantly between the three groups of infants (Supplemental Table 2 and Supplemental Figure 2). Notably, colonization by GBS CC17 was associated with increased abundance of three anaerobic genera, *i.e. Enterocloster*, *Finegoldia*, and *Veillonella*, compared to the other two cohort groups. Furthermore, colonization by non-CC17 GBS was associated with a decreased prevalence and abundance of *Enterobacter*, which are Gram-negative facultative anaerobes of the *Enterobacterales* order, compared to GBS CC17 colonized and GBS non-colonized neonates (p = 0.048 and p = 0.012, respectively).

**Figure 3.**
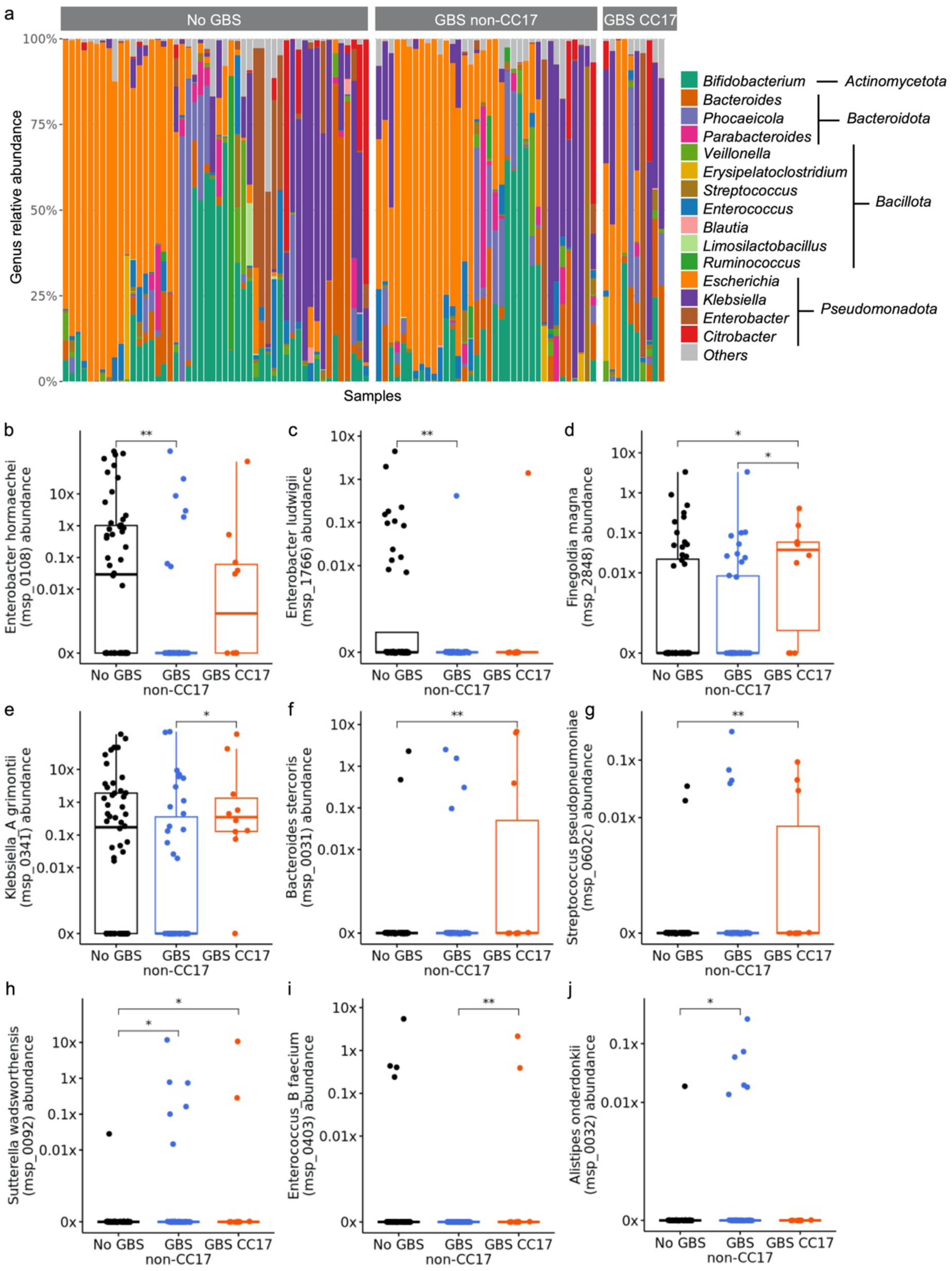
Metagenomics signatures of GBS colonization in fecal samples of 21 ± 7-day-old neonates. (a) Predominant genera with a relative abundance > 1% in the 100 fecal samples analyzed by shotgun metagenomics. (b-j) Abundances of the nine discriminant MGS according to GBS colonization: *Enterobacter hormaechei* (b), *Enterobacter ludwigii* (c), *Finegoldia magna* (d), *Klebsiella grimontii* (e), *Bacteroides stercoris* (f), *Streptococcus pseudopneumoniae* (g), *Sutterella wadsworthensis* (h), *Enterococcus faecium* (i), and *Alistipes onderdonkii* (j). (b-j) Data are displayed as box-and-whisker plots representing the interquartile range, with the central line indicating the median; individual data points are shown as dots. Differences in MGS abundance were determined using the Kruskal-Wallis test followed by the Wilcoxon rank sum test. * p<0.05, ** p<0.01. CC: clonal complex; GBS: Group B *Streptococcus*; MGS: metagenomic species.

To further characterize taxonomic signatures associated with GBS neonatal colonization, we compared MGS abundances across groups and identified 10 differentially abundant species (Figure 3b-j and Supplemental Table 3). As expected, the MGS representative of GBS (msp_2885) was the most contrasted (p = 4.0×10^-10^). The second most differentially abundant species was *Enterobacter hormaechei* (msp_0108, p = 0.0065, Figure 3b), which was significantly less abundant in GBS non-CC17 colonized neonates compared to GBS non-colonized neonates (p = 0.0019). Intriguingly, this association was not observed between GBS non-colonized and GBS CC17 colonized neonates (p = 0.45). A similar pattern was observed for another *Enterobacter* species, namely *Enterobacter ludwigii* (msp_1766, p=0.017, Figure 3c).

In addition, five MGS showed increased abundances in the fecal samples of GBS CC17 colonized neonates compared to GBS non-colonized and/or to GBS non-CC17 colonized neonates, including *Finegoldia magna* (msp_2848, p=0.037), *Klebsiella grimontii* (msp_0341, p=0.047), *Bacteroides stercoris* (msp_0031, p=0.023), *Streptococcus pseudopneumoniae* (msp_0602c, p=0.025), *Sutterella wadsworthensis* (msp_0092, p=0.031), and *Enterococcus faecium* (msp_0,403, p=0.048) (Figure 3d-i). Only one MGS, *Alistipes onderdonkii* (msp_0032), specifically showed an increased abundance in GBS non-CC17 colonized neonates compared to GBS non-colonized neonates (p=0.026, Figure 3j). Taken together, these findings suggest that neonatal colonization with GBS is associated with lineage-dependent shifts in specific bacterial taxa, without major differences in overall community structure.

### Formula-feeding contributes but does not fully explain differences in *E. hormaechei* abundance associated with GBS colonization

Given the importance of neonatal diet and mode of delivery in shaping the gut microbiota (Figure 2e-f) and the slight, albeit non-significant, association between formula feeding and GBS colonization status (Table 1), we evaluated their potential role as confounding factors in the observed association between GBS non-CC17 colonization and *E. hormaechei* abundance. We found that the prevalence and abundance of the *E. hormaechei* MGS were strongly associated with neonatal diet (Supplemental Figure 3a-b), being higher in mixed-fed and exclusively formula-fed neonates compared to those exclusively breastfed (p=0.019 and 0.0008 for the abundance comparison, respectively). Conversely, the mode of delivery was not associated with *E. hormaechei* or GBS abundance in our cohort (data not shown).

We next assessed whether diet alone could explain the association between *E. hormaechei* abundance and GBS colonization. A multivariable linear model including both diet and GBS status provided a significantly better fit than a model including diet alone (p = 8.0 × 10^-3^, ANOVA), indicating that GBS colonization contributes additional explanatory value beyond diet. Consistently, other taxa enriched in formula-fed infants, such as *Veillonella parvula* and *Klebsiella michiganensis*, showed no association with GBS colonization (p = 0.37 and 0.30, respectively). Finally, formula feeding was associated with increased microbiota diversity (Supplemental Figure 3c) as previously reported,^18^ whereas no differences were observed according to GBS colonization status (Figures 2b–c). Collectively, these findings indicate that while neonatal diet influences *E. hormaechei* abundance, it does not fully account for the differences observed between GBS non-CC17 colonized and GBS uncolonized neonates.

### Strain-level metagenomic analysis identifies the neonatal *E. hormaechei* isolates as *E. hormaechei* subsp. *hoffmannii* and *E. hormaechei* subsp. *steigerwaltii*

To gain insight into the interaction between GBS and *E. hormaechei*, we sought to identify the *E. hormaechei* strains present in our cohort at subspecies resolution. *E. hormaechei* is a member of the *Enterobacter cloacae* complex and comprises five closely related subspecies which can be distinguished by molecular approaches, including whole genome or *hsp60* amplicon sequencing.^20,21^ Using conventional culture and identification methods, we isolated colonies belonging to the *Enterobacter cloacae* complex from six neonatal fecal samples, in which *E. hormaechei* abundance by metagenomic analysis was the highest. Further *hsp60* sequencing allowed their identification as *E. hormaechei* subsp. *hoffmannii* (Ehh, n=4) and *E. hormaechei* subsp. *steigerwaltii* (Ehs, n=2).

To extend subspecies-level characterization across the entire cohort, we performed strain-level metagenomic profiling using PanPhlan. This analysis confirmed the results of *hsp60* sequencing and allowed the identification of *E. hormaechei* at the subspecies level in six additional fecal samples (Figure 4a). Although *E. hormaechei* was detected by metagenomics in 39 fecal samples, reliable strain-level profiles could not be obtained for 27 samples due to insufficient sequencing depth or evidence of strain mixtures. Overall, all identified *E. hormaechei* strains belonged either to Ehh (n=8) or Ehs (n=4).

**Figure 4.**
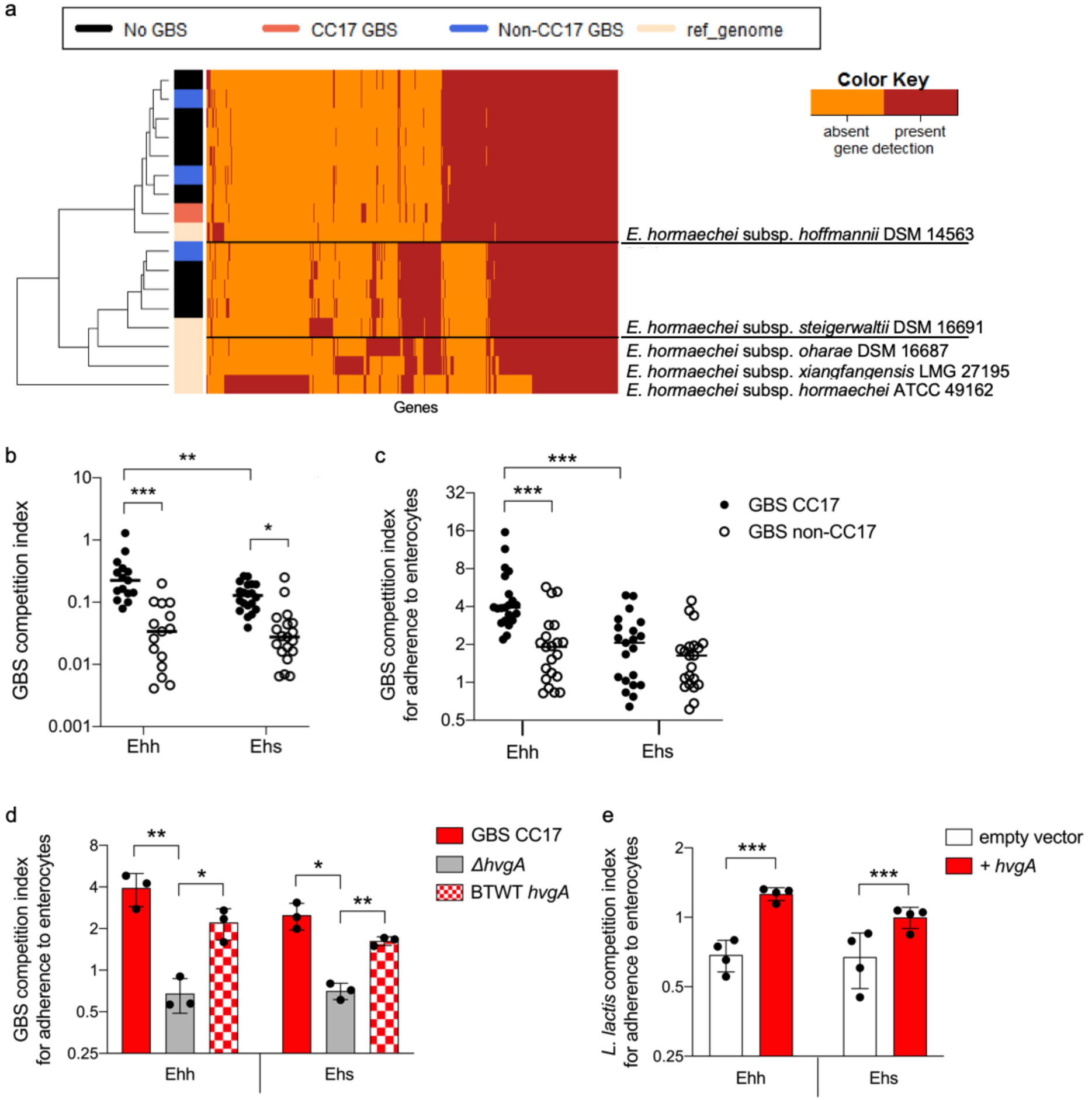
GBS CC17 better outcompetes *E. hormaechei* than GBS non-CC17. (a) Identification of *E. hormaechei* subspecies in 12 neonatal stool samples using strain-level analysis of metagenomics data. (b) GBS competition index against *E. hormaechei* in co-culture experiments in broth medium. Bacterial strains (4 Ehh, 4 Ehs, 5 GBS CC17 and 5 GBS non-CC17) were grown in Todd-Hewitt broth in co-culture (ratio GBS : *E. hormaechei* 1:1) and bacterial CFUs of each strain were enumerated following 16h of co-culture. Every GBS strain was tested against every *E. hormaechei* strain and experiments were performed twice in duplicate. (c) GBS competition index against *E. hormaechei* in co-adherence experiments on Caco-2 enterocytes. Bacterial strains (1 Ehh, 1 Ehs, 2 GBS CC17 and 2 GBS non-CC17) were tested for adherence in competition (ratio GBS : *E. hormaechei* 1:1) and bacterial CFUs of each strain were enumerated following 2h of adhesion at 37°C. Every GBS strain was tested against every *E. hormaechei* strain. Experiments were performed three times in duplicate. (b-c) Each dot represents the mean competition index for a given GBS - *E. hormaechei* pair with line at median. (d-e) Role of HvgA in GBS CC17 competition against *E. hormaechei* for adherence to enterocytes. Competition index of the Δ*hvgA* and the back-to-wild-type (BTWT) reverted mutant (d) and of *L. lactis* expressing HvgA (e) against *E. hormaechei*. (d-e) Results are expressed as mean +/- SD. Experiments were performed at least three times in triplicate. Dots represent the mean of triplicate values from individual experiments. (b to e) Competition indexes are expressed as the final ratio divided by the initial ratio between GBS or *L. lactis* and *E. hormaechei* CFUs. (b-e) Statistical analyses were performed by 2-way ANOVA. * p<0.01; ** p<0.01; *** p<0.001. CC: clonal complex; Ehh: *E. hormaechei* subsp. *hoffmannii*; Ehs: *E. hormaechei* subs. *steigerwaltii*; GBS: Group B *Streptococcus*. SD: standard deviation.

### GBS CC17 strains display increased competitive fitness against *E. hormaechei* in co-culture

To investigate the GBS-*E. hormaechei* interactions underlying the associations identified by metagenomics, and particularly the lower abundance of *E. hormaechei* in fecal samples of GBS non-CC17 carriers, we first assessed whether direct antagonistic activity could occur using an overlay screening assay.^22^ No bacteriocin-like activity between the two species was detected (data not shown).

Next, we evaluated interspecies competition in co-culture experiments in liquid medium, using human clinical isolates of Ehh (n=4), Ehs (n=4), GBS CC17 (n=5), and GBS non-CC17 (n=5) (Supplemental Table 1). Both species were able to grow in co-culture, reaching final densities of approx. 1.10^9^ and 1.10^8^ colony forming units (CFU)/mL, respectively (Supplemental Figure 4a), indicating coexistence despite competition. Nevertheless, *E. hormaechei* consistently outcompeted GBS strains, with a significantly stronger effect on non-CC17 than on CC17 strains (Figure 4b). Indeed, the final GBS densities in co-culture with *E. hormaechei* were significantly higher for GBS CC17 than for non-CC17 strains (approximately 1.7x10^8^ and 1.0x10^8^ CFU/mL, respectively, p<0.001, Supplemental Figure 4b), indicating increased competitive fitness of the CC17 lineage.

To determine whether intrinsic growth properties could account for this phenotype, we compared the growth characteristics of GBS CC17 and non-CC17 strains in monoculture. Lag phase duration (Mean ± SD = 73 ± 26 and 142 ± 34 min, respectively) and doubling time (Mean ± SD = 50 ± 9 and 45 ± 3 min, respectively) were similar and could not explain the competitive advantage of GBS CC17 (Supplemental Figure 5).

### GBS CC17 strains display enhanced competitive fitness against *E. hormaechei* for adherence to enterocytes

The absence of bacteriocin-like activity between GBS and *E. hormaechei*, together with the ability of all tested strains to grow in co-culture, suggested that the lower abundance of *E. hormaechei* in fecal samples of GBS non-CC17 carriers could be due to an impaired capacity of non-CC17 GBS isolates to compete with *E. hormaechei* for gut colonization, rather than to the inability of *E. hormaechei* to colonize the gut in presence of non-CC17 GBS. Given that such negative association was not observed for GBS CC17, and that GBS CC17 strains were previously reported to exhibit increased adherence to enterocytes compared to non-CC17 strains,^9^ we hypothesized that GBS CC17 isolates could better compete against *E. hormaechei* for gut colonization and especially for adherence to enterocytes.

To test this hypothesis, we first measured the adherence of three Ehh and three Ehs isolates to Caco-2 enterocytes and found similar adherence capacities, allowing the selection of one representative strain of each subspecies for competition experiments (Supplemental Figure 6a). Using two representative GBS CC17 and non-CC17 isolates, including the reference strains BM110 (CC17) and NEM316 (CC23) (Supplemental Figure 6b), we found that GBS CC17 adherence to enterocytes was neither affected by Ehh nor by Ehs, whereas that of non-CC17 GBS was inhibited 2-fold by Ehs (Supplemental Figure 6c). Besides, GBS CC17 reduced the adherence of both Ehh and Ehs to enterocytes, whereas non-CC17 GBS only inhibited Ehs adherence (Supplemental Figure 6d). Eventually, both GBS CC17 and non-CC17 outcompeted Ehh for adherence to enterocytes, GBS CC17 being twice more effective than non-CC17 GBS (median competition indexes of 4 and 2, respectively, Figure 4c). In contrast, the ability of GBS CC17 isolates to outcompete Ehs was weaker (median competition index of 2, range 0.5 – 5) and was similar to that of non-CC17 strains.

### HvgA mediates the competitive advantage of GBS CC17 against *E. hormaechei* for adherence to enterocytes

The enhanced adherence of GBS CC17 to enterocytes has previously been attributed to the expression of the CC17-specific surface protein HvgA which confers increased adherent properties to multiple cell types.^9^ Therefore, we investigated whether HvgA could contribute to the competitive advantage of GBS CC17 over *E. hormaechei* for adherence to enterocytes. As shown in Figure 4d, the ability of a Δ*hvgA* mutant to compete with Ehh or Ehs for cell adherence was abolished (competition index < 1). GBS competitive advantage was restored in the reverted mutant (back-to-wild-type), suggesting a direct contribution of HvgA. To further assess whether HvgA expression was sufficient to confer enhanced competitive fitness, we performed competition experiments using *Lactococcus lactis*, a Gram-positive non-pathogenic species closely related to GBS with poor adherent properties. The heterologous expression of HvgA in *L. lactis*^9^ was sufficient to increase its competition capacities for adherence to enterocytes against Ehh and Ehs (Figure 4e). Together, these findings identify HvgA as a key determinant of the enhanced competition for adherence to enterocytes displayed by GBS CC17.

## Discussion

GBS is recognized as the leading cause of neonatal invasive infections worldwide, with maternal recto-vaginal colonization being the major risk factor for both early- and late-onset diseases.^16,23,24^ Strategies based on intrapartum antibiotic prophylaxis (IAP) for colonized women have proven to be efficient in preventing vertical transmission and neonatal early-onset sepsis, whose incidence has decreased by 80% in the past decades.^6,16,25^ Conversely, the incidence of GBS late-onset disease has remained stable and now exceeds that of early-onset disease in many countries, underscoring the necessity for alternative preventive strategies.^10,11,13^ While the exact source and mechanisms of neonatal contamination are still unclear, mounting evidence suggests that the gut may be the primary portal of entry for late-onset GBS disease.^7,9,26^ This hypothesis is supported by observations that GBS can be detected in infant stool prior to late-onset infection.^7,26^ Therefore, a more thorough understanding of the mechanisms that lead to GBS establishment and persistence in the neonatal gut could result in the development of innovative preventive strategies against late-onset infections.

In this study, we used shotgun metagenomic sequencing with a specific gene catalogue of the infant gut to deeply analyze, at species-level resolution, the fecal microbiota of 21-day-old neonates born at term to GBS-colonized mothers. We identified specific taxonomic signatures of neonatal GBS colonization including decreased abundance of *E. hormaechei* in neonates colonized with GBS non-CC17 strains compared to GBS-uncolonized neonates. Focusing on the hypervirulent GBS CC17, which is responsible for approximately 80% of late-onset neonatal cases,^11,24^ we further identified distinct microbiota signatures associated with CC17 colonization, including enrichment in taxa such as *F. magna* and *E. faecium*. Notably, the lower abundance of *E. hormaechei* was not observed in CC17-colonized neonates. Our experimental data suggest that this lineage-specific pattern may result from enhanced competitive fitness of GBS CC17 for gut colonization, mediated at least in part by the CC17-specific HvgA adhesin.

In infants, gut microbiota establishment and composition is primarily shaped by the mode of delivery, the gestational age at delivery, the neonatal feeding diet, and also by IAP.^27–29^ Using 16S rRNA gene sequencing, a few studies investigated the impact of GBS maternal colonization itself on neonatal fecal microbiota up to 6 months of age.^15,30,31^ Overall, these studies did not identify major differences in fecal microbiota α or ß diversity between infants born to GBS-negative and GBS-positive mothers. However, differences in abundance of specific taxa, including *Ruminococcus*, *Clostridium*, *Akkermansia*, and *Bacteroides*, have been reported, even when adjusting for IAP.^15^ Consistent with these observations, GBS colonization in our cohort did not significantly impact microbiota α diversity and overall community structure. Altogether, these findings suggest that GBS is not a major driver of gut microbiota composition in infants. Nevertheless, GBS colonization is associated with differential abundances of specific bacterial species that vary depending on the GBS lineage, suggesting the existence of specific interspecies interactions, whether direct or indirect.

The majority of GBS-microbiota interactions that have been studied so far were in the context of maternal vaginal colonization.^32,33^ Both synergistic and antagonistic interactions with diverse species including *Lactobacillus* spp., *Staphylococcus aureus*, and *Candida albicans* have been described, both in clinical and experimental models. However, extrapolation of these findings to the neonatal gut remains challenging given the dynamic nature of the developing infant microbiota, dominated by pioneer colonizers such as *Bifidobacteriales* and *Enterobacterales.*^34^ Despite the marked differences between the vaginal and gut microbiomes, some overlapping ecological associations have nevertheless been reported. Consistent with our findings, specific taxa including the genus *Veillonella* and *F. magna* were found to be positively correlated with vaginal GBS colonization by 16S rRNA gene sequencing.^35^ Interestingly, *Veillonella* are saccharolytic bacteria which use lactate, the end product of lactic acid bacteria including *Streptococcus* spp., to produce propionate. *Veillonella* and *Streptococcus* are frequently described as co-occurring.^36^ However, the mechanisms underlying the specific association between GBS CC17 colonization and the abundance of *Veillonella* in our cohort remain unclear.

Bacteriocins are key elements in interspecies competition for niche colonization. However, we did not observe any bacteriocin-like activity between GBS and *E. hormaechei*. In contrast to many streptococcal species, little is known about bacteriocin production in GBS. In a screen for bacteriocin-like activity among GBS isolates, only 5% exhibited an inhibitory activity against other bacterial species.^37^ More recently, two GBS bacteriocins designated Agalacticin and Nisin P were described.^38–40^ Both peptides exhibit inhibitory activity against Gram-positive bacteria but not against Gram-negatives such as *Pseudomonas* spp. and *E. coli*. Neither peptide appeared to be highly conserved among GBS isolates and the significance of bacteriocin production in GBS capacity to colonize diverse host niches remains to be elucidated. In Gram-positive bacteria, interspecies competition can also be mediated by type VII secretion systems (T7SS).^41^ T7SS secrete effector proteins with functions in virulence, host cell toxicity, and interbacterial killing. In GBS, four distinct subtypes of T7SS have been identified.^42^ Though imperfect, a correlation exists between the encoded T7SS subtype and the various GBS lineages.^42,43^ Interestingly, GBS CC17 isolates specifically encode the T7SS subtype IV, which is remarkably the only subtype that does not encode effector proteins. Consequently, it is plausible that the T7SS is not functional in this particular lineage, and that the colonization fitness of GBS CC17 do not rely on direct interspecies competition but rather on its intrinsic capacity to interact with host cells. The absence of a functional T7SS might also contribute to the lower level of intestinal colonization of GBS CC17 strains compared to non-CC17 strains.

Consistent with this hypothesis, both GBS CC17 and non-CC17 strains outcompeted *E. hormaechei* for adherence to enterocytes in our experiments, although CC17 strains were even more effective against Ehh isolates. These findings suggest that *E. hormaechei* gut colonization might inhibit GBS establishment in the neonatal gut, an effect that could be partially mitigated for GBS CC17 strains due to their enhanced adherence capacities to enterocytes. Importantly, we found that *E. hormaechei* abundance and the introduction of formula feeding were both associated to increased microbiota α diversity, a finding which suggests that microbiota diversification following infants’ diet diversification contributes to lowering GBS colonization. This effect may, however, be attenuated in the case of the hypervirulent GBS CC17, possibly because of its enhanced capacity to colonize the neonatal gut.^9,14^

Our study took advantage of a well-characterized and highly homogenous cohort regarding gestational age, mode of delivery, IAP, and feeding diet, all of which being major drivers of microbiota composition in early life.^34^ This design reduced, although did not eliminate, potential confounding factors and allowed us to specifically investigate the influence of gut microbiota on GBS colonization in the first month of life. The negative association observed between *E. hormaechei* and GBS non-CC17 was further supported by *in vitro* competition experiments, strengthening the biological plausibility of our metagenomic findings. However, other associations, including the co-occurrence of GBS CC17 with several taxa, remain to be functionally elucidated. In addition, our conclusions are limited by the number of GBS CC17 colonized neonates analyzed in this study. Because of the limited size of our cohort, other confounding variables, such as ethnicity or maternal diet could not be investigated. Additionally, other associations between microbiota composition and GBS colonization status might have been missed.

In conclusion, we identified specific and distinct microbial signatures of neonatal colonization by the hypervirulent GBS CC17 and by non-CC17 GBS, suggesting different interspecies and host-microbiota interactions that could account for the overrepresentation of GBS CC17 in neonatal late-onset infections. Although we did not observe any association between GBS colonization and typical probiotic taxa, such as lactobacilli and bifidobacteria, this does not exclude a potential role for specific beneficial microbes in modulating GBS colonization. These findings support further research on microbiota-based strategies aimed at preventing neonatal colonization and infection by pathobionts.

## Methods

### Study design and participants

Metagenomics was performed on a subset of fecal samples collected through the prospective Col-StreptoB cohort study conducted in France between November 2012 and April 2015 (ClinicalTrials.gov #NCT01719510). The study aimed at identifying demographic, clinical, and risk factors for GBS colonization in neonates and infants. The design and results have been published and are briefly summarized below.^14^ The eligibility criteria for maternal participation in the study included all pregnant women over 18 years-old who were screened positive for vaginal GBS colonization between 34-38 weeks of gestational age and women with a pregnancy over 34 weeks of gestational age screened positive at time of delivery. Women positive for GBS colonization were administered penicillin G as the first line antibiotic for intrapartum antibiotic prophylaxis (IAP), and cefazolin or clindamycin as second line antibiotics in case of mild or severe allergy to penicillin, respectively, as recommended by the French guidelines for the prevention of GBS early-onset neonatal sepsis.^16,17^

Mother-infant pairs were recruited and monitored after maternity discharge at 21 ± 7 days and 60 ± 7 days. Demographic and clinical data (maternal age, maternal place of birth, pregnancy characteristics such as parity and pregnancy-associated complications, delivery characteristics such as the mode of delivery and the duration of IAP, infant antibiotic therapy, infant diet) were collected in a computerized database. At discharge from the maternity, all women received the material for oral swabbing and neonatal stool sampling. Stools were sampled in the diapers by swabs collected in Amies transport medium (ESwab™ system, Copan Diagnostics, Italy). Samples were stored at 4°C before being sent to the laboratory within 48h where they were kept at 4°C before being processed within 48h for the detection of GBS using conventional culture methods and stored at -80°C for further analyses. A molecular typing for the identification of GBS CC17 based on the detection of the *hvgA* gene was performed for all GBS isolates.^44^

Eventually, the Col-StreptoB study included 890 mother-infant pairs. At 21 ± 7 days old, 748 dyads completed follow-up and 157 neonates (21%) were found colonized by GBS. Neonatal and maternal isolates were of the same capsular type and CC in 95% of the cases, pointing at the mother as the most likely source of neonatal contamination. Metagenomic analysis was performed on a selection of 50 GBS-positive and 52 GBS-negative fecal samples (Figure 1). This selection included all the samples positive for GBS CC17 (n=10). For the other samples, a random selection was made to obtain approximately half breastfed and half formula-fed neonates in each group.

### Ethical Approval and Informed Consent statement

This study was conducted in accordance with the Helsinki Declaration, in agreement with French regulations on privacy, data collection and treatment, and was approved by the Comité consultatif sur le traitement de l’information en matière de recherche (CCTIRS, Advisory committee on information processing for research, authorization 12005). The CoStreptoB study was registered at ClinicalTrials.gov, number NCT01719510. All mothers included and the respective fathers gave written informed consent.

### DNA extraction and shotgun metagenomic sequencing

Fecal DNA was extracted following the SOP 07 V2 H from IHMS.^45,46^ The DNA preparation was subjected to quality control using Qubit Fluorometric (ThermoFisher Scientific, Waltham, US) and qualified using DNA size profiling on a Fragment Analyzer instrument (Agilent Technologies, Santa Clara, US). A total of 3 µg of high molecular weight DNA (>10 kbp) was used to build sequencing libraries. Shearing of DNA into fragments of approximately 150 bp was performed using an ultrasonicator (Covaris, Woburn, US) and DNA fragment library construction was performed using the Ion Plus Fragment Library and Ion Xpress Barcode Adapters Kits (ThermoFisher Scientific, Waltham, US). Purified and amplified DNA fragment libraries were sequenced using the Ion Proton Sequencer (ThermoFisher Scientific, Waltham, US).

### Sequencing data preprocessing

First, quality control was performed with AlienTrimmer:^47^ 1) sequencing adapters were removed, 2) low quality reads were trimmed or discarded, and 3) too short reads (< 60 bp) were discarded. Then, reads mapped to the human genome (T2T CHM13v2.0 GCA_009914755.4) with bowtie2^48^ were removed. Finally, 11M high-quality reads were randomly selected in each sample with fastq-sample.^49^

### Gene abundance table generation

The gene abundance table was generated with the METEOR software.^50^ First, selected high quality reads were mapped with bowtie2^48^ to a gene catalog of the infant gut microbiota, comprising 2.4 millions of genes. Alignments with nucleotide identity < 95% were discarded and gene counts were computed with a two-step procedure previously described that handles multi-mapped reads.^51^ Finally, raw gene counts were normalized according to gene length.

### Creation and annotation of MetaGenomic Species (MGS)

Co-abundant genes across samples from this study and publicly available cohorts^18,19^ were binned in MGS Pan-genomes using MSPminer.^52^ Quality control of each MGS was manually performed by visualizing heatmaps representative of the normalized gene abundance profiles. In addition, MGS completeness and contamination was assessed by searching for 40 universal single copy marker genes with fetchMGs^53^ and by checking taxonomic homogeneity. Taxonomic annotation of the MGS was carried out with GTDB-Tk^54^ based on GTDB Release 05-RS95.

### MGS abundance table generation

The abundance of an MGS in a sample was defined as the mean abundance of its 100 marker genes, *e.g.* species-specific core genes that correlate the most altogether. If less than 10% of the marker genes were seen in a sample, the abundance of the MGS was considered as null. Abundances at higher taxonomic ranks were computed as the sum of the MGS that belong to a given taxa.

### Statistical analysis of metagenomic data

All statistical analyses were conducted using R (version 4.4.1) primarily using packages from the tidyverse suite for data processing and visualization. For α diversity, species richness was defined as the number of MGS with strictly positive abundance in a sample, and the Shannon index was computed using the diversity function from the vegan package (version: 2.6-6.1). To assess differential abundance, the Mann-Whitney U test was employed for pairwise comparisons, and the Kruskal-Wallis test was used for comparisons involving three groups. Differences in taxa prevalence (presence/absence) were determined using Fisher’s exact test for two-group comparisons and the Chi-square test (χ2) for multiple-group comparisons. Multiple-testing correction was performed.

Beta diversity was assessed using the Bray-Curtis dissimilarity computed on the log10-transformed MGS abundance table with the vegan package. Principal Coordinates Analysis (PCoA) was performed using the pcoa function from the ape package (version 5.8). Group differences in community structure were tested using Permutational Multivariate Analysis of Variance (PERMANOVA), performed with the adonis function (permutations = 9,999) from the vegan package.

### Detection of GBS surface proteins encoding genes for *in silico* typing

The identification of GBS CC17 and GBS non-CC17 in sequencing reads was based on the detection of the *srr2* and *hvgA* CC17-specific surface proteins encoding genes and on their non-CC17 respective homologues *srr1* and *bibA.*^44,55^ The genes *srr1* and *bibA* found in the genome of GBS strain NEM316^56^ were downloaded from GenBank (accession numbers CAD47188.1 and CAD47677.1, respectively). Similarly, the CC-17-specific genes *srr2* and *hvgA* found in the genome of GBS strain BM110^57^ were downloaded (accession numbers SIW57866.1 and SIW58571.1, respectively). For *srr2*, only the repeats-free binding region (BR2) which goes from bases 574 to 1656 was considered. For *srr1*, only the homologous BR1 region going from bases 350 à 1700 was considered. For *bibA* and *hvgA,* the first 100 nucleotides which correspond to a secretion signal sequence, as well as the last 105 nucleotides which correspond to the anchoring signal in the bacterial cell wall were filtered out.

Sequencing reads were mapped on the specific regions of the four genes described above with bowtie2^48^ (parameters: --local-sensitive). End-to-end alignments with nucleotide identity ≥ 95% were considered. Local alignments were considered only if they started or ended at the edge of one of the regions of interest.

### Strain-level metagenomic analysis of *Enterobacter hormaechei* subspecies

Subspecies of *E. hormaechei* were identified in metagenomic samples with PanPhlan version 1.2.3.6.^58^ First, a pangenome was built using the script “pangenome_generation.py” with all the coding DNA sequences (CDS) predicted in *E. hormaechei* subsp*. hoffmannii* DSM 14563 (accession: GCA_001729745.1), *E. hormaechei* subsp*. steigerwaltii* DSM 16691 (accession: GCA_001729725.1), *E. hormaechei* subsp*. oharae* DSM 16687 (accession: GCA_001729705.1), *E. hormaechei* subsp*. xiangfangensis* LMG27195 (accession: GCA_001729785.1) and *E. hormaechei* subsp*. hormaechei* ATCC 49162 (accession: GCA_001875655.1). Then, sequencing reads of each sample were mapped on the *E. hormaechei* pangenome with the script “panphlan_map.py”. Finally, a gene presence/absence table including metagenomic samples and reference genomes was generated with the “panphlan_profile.py” script. Notably, samples where the presence of multiple closely related strains was suspected or where the strain had low sequencing coverage (< 3x) were excluded. For visualization purposes, complete linkage clustering of samples in rows and genes in columns was performed using the Jaccard distance.

### Bacteria and cell lines

Bacterial strains used in this study are listed in Supplemental Table 1. GBS strain BM110, capsular type III, multi-locus sequence type (ST) 17 and CC17, and GBS strain NEM316, capsular type III, ST23 and CC23, are well characterized isolates from humans with invasive infections.^56,59^ Other GBS and *E. hormaechei* strains were isolated from human samples. GBS BM110 in-frame deletion mutant of *hvgA* was constructed by double crossing-over with a deleted copy of the gene as previously described,^9^ leading to the selection of the mutant and the back-to-wild-type complemented strain. *L. lactis* expressing HvgA was previously constructed using the pOri23 shuttle vector.^9^ GBS and *L. lactis* were cultured at 37°C and 30°C, respectively, in Todd Hewitt broth or agar, and erythromycin was used at the concentration of 10 µg/mL for vector-containing *L. lactis*. Unless otherwise specified, *E. hormaechei* isolates were cultured at 37°C in Trypticase soja broth or agar. Growth curves and doubling times of GBS and *E. hormaechei* clinical isolates were obtained by measuring the optical density at 600 nm (OD_600nm_) every 10 minutes for 12 hours using a Multiskan^TM^ GO microplate spectrophotometer (ThermoFisher).

The human colonic epithelial cell line Caco-2/TC7^60^ (RRID:CVCL_0233) was cultured at 37°C in a 10% CO_2_ atmosphere in DMEM high glucose with GlutaMax (ThermoFisher) supplemented with pyruvate (ThermoFisher) and non-essential amino-acids (ThermoFisher) with 20% inactivated fecal calf serum (FCS, Eurobio). Cells were tested for mycoplasma using the MycoAlert mycoplasma detection kit (Lonza) every 2 to 3 weeks and always found to be negative.

### Bacterial co-culture experiments

For GBS and *E. hormaechei* growth experiments in co-culture, bacteria in mid-exponential growth phase (OD_600nm_=0.5) were harvested by centrifugation (10 min at 4,000 g), washed twice with Phosphate Buffer Saline (PBS) buffer, and re-suspended in PBS buffer. Todd-Hewitt broth (10 mL) was inoculated with a 1:1 ratio of GBS and *E. hormaechei* at a total density of 5.10^6^ CFU/mL. Bacteria were enumerated following 16h of incubation at 37°C without agitation.

### Adherence assays

Caco-2 cells were seeded on standard Costar plates (ThermoFisher) at a density of 2x10^5^/cm^2^ and incubated 48h to 72h until reaching confluency before being washed twice in PBS and infected by the bacteria. Bacteria in mid-exponential growth phase (OD_600nm_=0.5) were harvested by centrifugation (10 min at 4,000 g), washed twice with PBS buffer, and re-suspended in cellular growth medium without FCS. Cells were infected at a multiplicity of infection (moi) of 10 bacteria per cell. For competition experiments, cells were infected with a 1:1 ratio of GBS or *L. lactis* and *E. hormaechei* at a total moi of 10 bacteria per cell. Following 2h of incubation at 37°C in a 10% CO_2_ atmosphere, cells were washed 5 times with PBS and cell lysates were recovered for quantification of adherent bacteria.

### Co-culture counts and competition index determination

In co-culture and co-adherence experiments, serial dilutions of growth medium or cell lysates were plated on trypticase soja agar for selective isolation of *E. hormaechei* and on TH agar supplemented with colistin (10 mg/L) for selective isolation of GBS or *L. lactis*. Competition indexes were calculated as the final ratio between GBS or *L. lactis* and *E. hormaechei* colony forming units (CFU) divided by the ratio between GBS or *L. lactis* and *E. hormaechei* CFU of the initial inoculum.

### Statistical analysis of *in vitro* experimental data

Statistical analysis was performed using GraphPad Prism 5.0 software. Multiple-group comparisons were performed by Kruskal-Wallis or two-way ANOVA. A p value < 0.05 was considered significant.

## Data availability statement

Metagenomic sequencing data have been deposited in the European Nucleotide Archive (ENA) under BioProject accession number PRJEB101714.

The other datasets generated during the current study are available from the corresponding author on reasonable request.

## Acknowledgements

The authors thank the management of the Col-StreptoB clinical trial Florence Artiguebieille, Valerie Fauroux, Farah Ketroussi, and Chahrazed Guettouche from Unité de recherche clinique - centre d’investigation clinique Cochin-Necker, Assistance Publique - Hôpitaux de Paris, Paris. They thank all the participants, mothers and fathers, as well as the clinical teams for their assistance in recruiting patients to the study.

## Authors contribution

Conceptualization: S.D.E., C.Po, A.T. Methodology: AS.A., F.P.O., G.T., S.K., F.G., L.M., S.D.E., C.Po., A.T. Resources: AS.A., F.P.O., F.G., C.Pl., L.M., S.D.E., C.Po. A.T. Data Curation: AS.A., F.P.O., S.K., C.Pl. A.T. Investigation: AS.A., F.P.O., G.T., S.K., A.T. Formal Analysis: AS.A., F.P.O., A.T. Software; AS.A., F.P.O. Visualization: AS.A., F.P.O., G.T., A.T. Supervision: S.K., S.D.E., C.Po., A.T. Funding Acquisition: F.G., S.D.E., C.Po. Writing – original draft: AS.A., F.P.O., A.T. Writing – review and editing: all authors.

## Competing interests

The authors report there are no competing interests to declare.

### Material and correspondence

Correspondence and material requests should be addressed to AT.

## Supplemental Material

**Supplemental Table 1.** Bacterial strains used in this study.

**Supplemental Table 2.** Statistical comparisons of discriminant genera between GBS uncolonized, GBS non-CC17 colonized, and GBS CC17 colonized neonates at 21 ± 7 days of age.

**Supplemental Table 3.** Statistical comparisons of discriminant metagenomic species between GBS uncolonized, GBS non-CC17 colonized, and GBS CC17 colonized neonates at 21± 7 days of age.

Supplemental Figure 1. E*s*cherichia *coli* abundance (a) and prevalence (b) in neonatal fecal samples.

Supplemental Figure 2. Discriminant genera between GBS uncolonized, GBS non-CC17 colonized, and GBS CC17 colonized neonates at 21± 7 days of age.

Supplemental Figure 3. Impact of neonatal diet on *E. hormaechei* abundance and microbiota diversity.

Supplemental Figure 4. *E. hormaechei* and Group B *Streptococcus* growth in co-culture in broth medium.

Supplemental Figure 5. *E. hormaechei* and Group B *Streptococcus* growth characteristics in broth medium.

Supplemental Figure 6. Adherence of *E. hormaechei* and Group B *Streptococcus* on Caco-2 enterocytes.

