## Supplemental Table 1 and Figures for "Competitive advantage of hypervirulent Group B *Streptococcus* for neonatal colonization revealed by metagenomic analysis of gut microbiota"

**Supplemental Table 1. Bacterial strains used in this study.**

| Species | Subspecies / subtype | Strain | Origin - Description | Source |
|---|---|---|---|---|
| *Enterobacter cloacace* | | | | |
| | subsp. *hoffmannii* | CCH 2064 | Neonatal stool sample (ColStreptoB trial) | This study |
| | subsp. *hoffmannii* | CCH 2065 | Neonatal stool sample (ColStreptoB trial) | This study |
| | subsp. *hoffmannii* | CCH 2066 | Neonatal stool sample (ColStreptoB trial) | This study |
| | subsp. *hoffmannii* | CCH 2067 | Neonatal stool sample (ColStreptoB trial) | This study |
| | subsp. *steigerwaltii* | CCH 2060 | Neonatal nasopharynx | [1] |
| | subsp. *steigerwaltii* | CCH 2061 | Adult urine sample | [1] |
| | subsp. *steigerwaltii* | CCH 2062 | Adult sputum sample | [1] |
| | subsp. *steigerwaltii* | CCH 2063 | Neonatal stool sample (ColStreptoB trial) | This study |
| *Streptococcus agalactiae* | | | | |
| | CC17 – CPS III | BM110 | Neonatal blood culture | [2] |
| | CC17 – CPS III | CNR CCH 2017-1486 | Neonatal cerebrospinal fluid, this study | This study |
| | CC17 – CPS III | CNR CCH 2017-1496 | Neonatal cerebrospinal fluid, this study | This study |
| | CC17 – CPS III | CNR CCH 2017-1510 | Adult blood culture, this study | This study |
| | CC17 – CPS III | CNR CCH 2017-1511 | Adult Neonatal blood culture, this study | This study |
| | CC23 – CPS III | NEM316 | Neonatal blood culture | [3] |
| | CC19 – CPS V | CNR CCH 2017-1676 | Neonatal gastric fluid, this study | This study |
| | CC19 – CPS III | CNR CCH 2017-1697 | Adult vaginal sample, this study | This study |
| | CC10 – CPS Ib | CNR CCH 2017-1700 | Adult blood culture, this study | This study |
| | CC23 – CPS Ia | CNR CCH 2017-1731 | Adult blood culture, this study | This study |
| | CC17 – CPS III | BM110 Δ*hvgA* | In frame deletion mutant | [4] |
| | CC17 – CPS III | BM110 Δ*hvgA* BTWT | Δ*hvgA* complemented strain | This study |
| *Lactococcus lactis* subsp. *cremoris* | | | | |
| | | MG1363 | | [5] |
| | | MG1363, pOri23 | *L. lactis* with empty vector | [4] |
| | | MG1363, pOri23Ω*hvgA* | *L. lactis* expressing HvgA | [4] |

BTWT: back-to-wild-type; CC: clonal complex; CPS: capsular type.

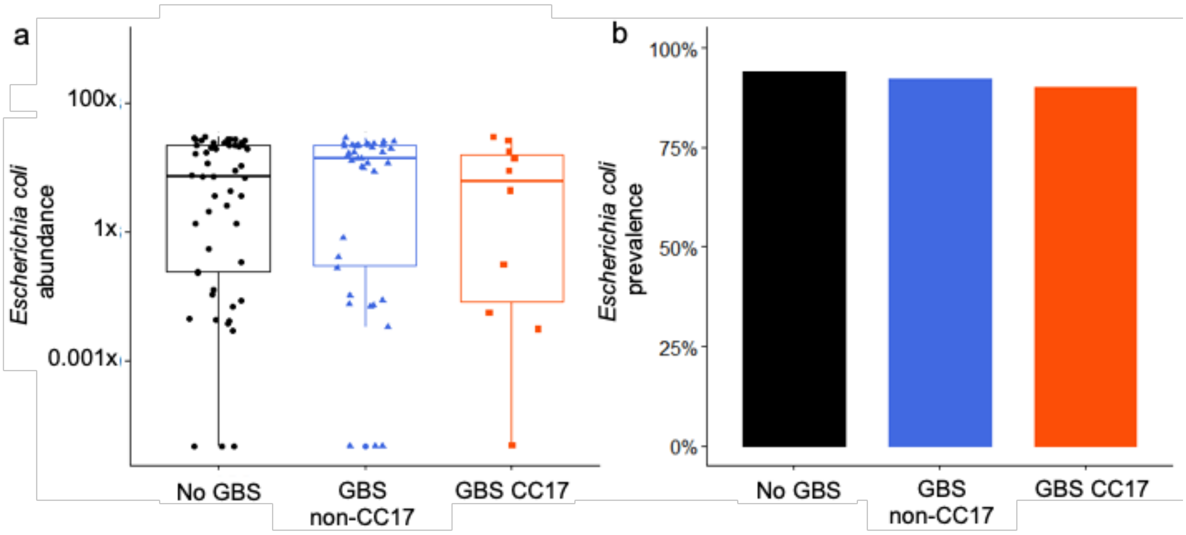


**Supplemental Figure 1. *Escherichia coli* abundance (a) and prevalence (b) in neonatal fecal samples.** (a) Data are displayed as box-and-whisker plots representing the interquartile range, with the central line indicating the median; individual data points are shown as dots. Statistical analyses were performed using the Kruskal Wallis test followed by the Wilcoxon rank sum test (a) and Fisher's exact test (b). CC: clonal complex; GBS: Group B *Streptococcus*.

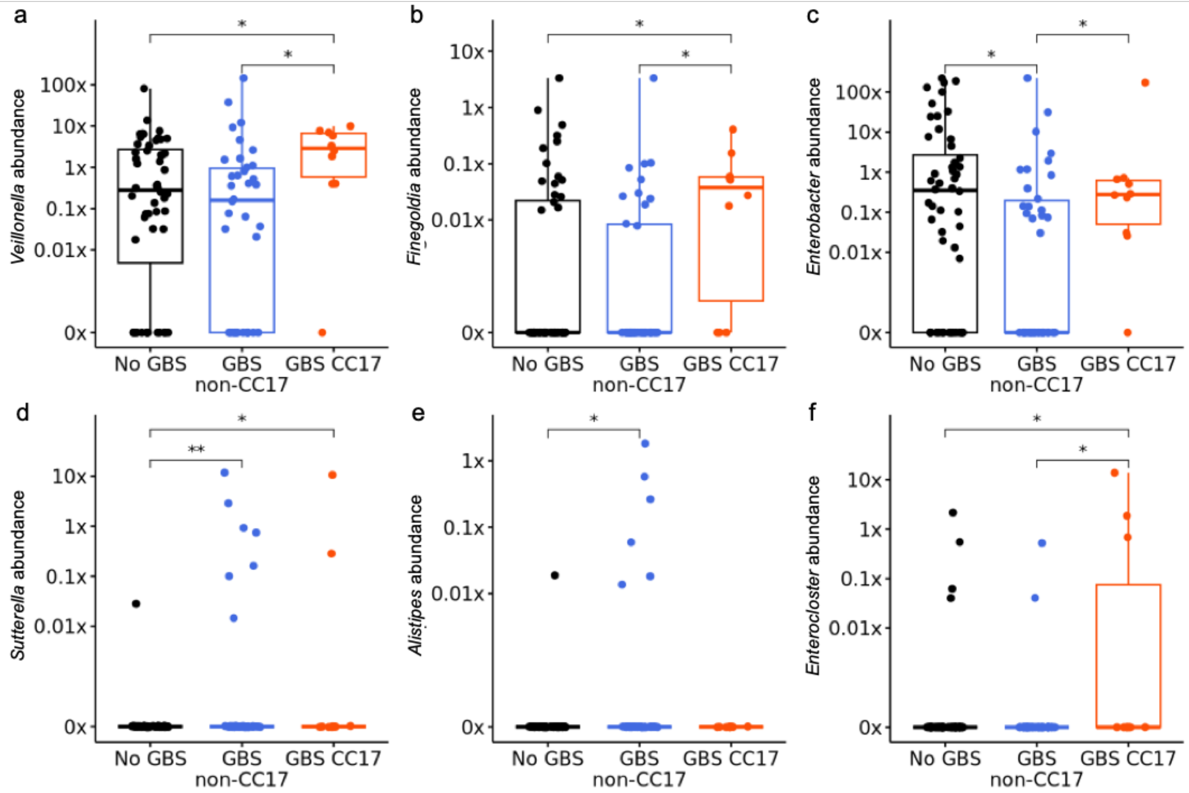


**Supplemental Figure 2. Discriminant genera between the three cohort groups.** Abundances of *Veillonella* spp. (a), *Finegoldia* spp. (b), *Enterobacter* spp. (c), *Sutterella* spp. (d), *Alistipes* spp. (e), and *Enterocloster* spp. (f). Data are displayed as box-and-whisker plots representing the interquartile range, with the central line indicating the median; individual data points are shown as dots. Differences in taxa abundance were determined using the Kruskal-Wallis test followed by the Wilcoxon rank sum test. * p<0.05, ** p<0.01. CC: clonal complex; GBS: Group B *Streptococcus*.

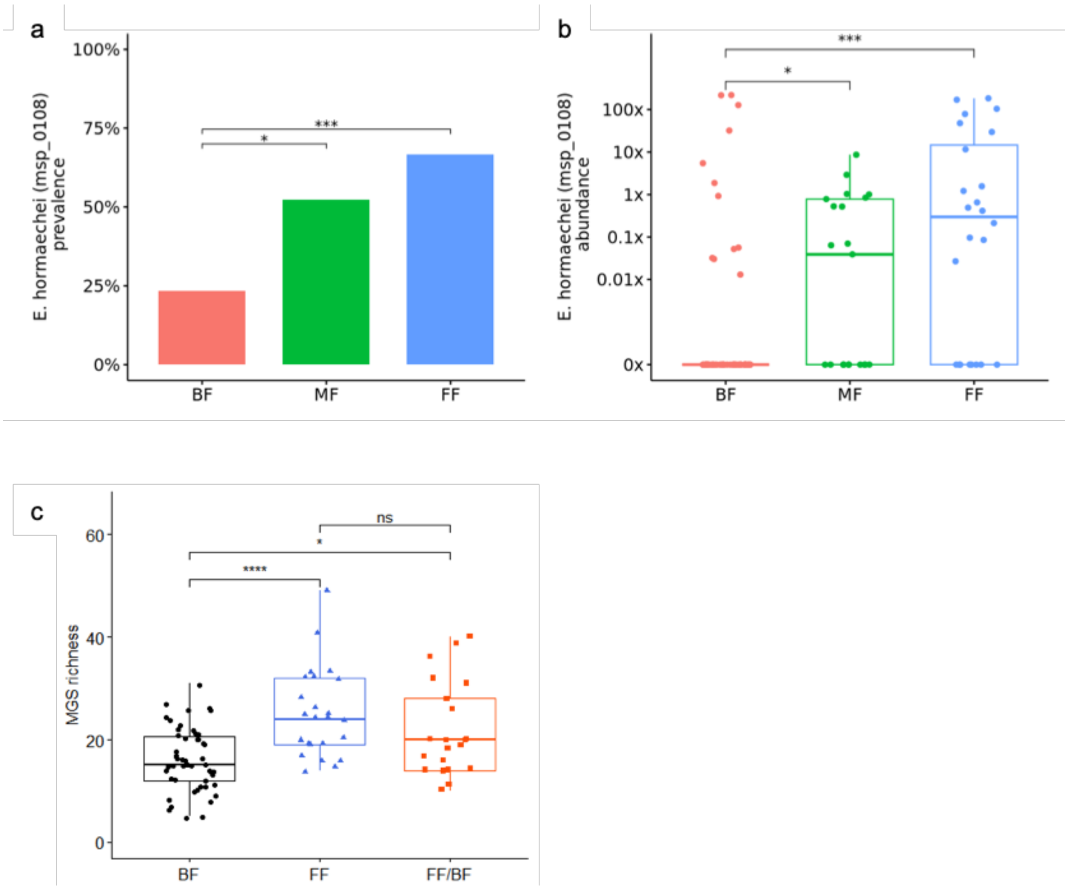


**Supplemental Figure 3. Impact of neonatal diet on *E. hormaechei* abundance and microbiota diversity.** *E. hormaechei* prevalence (a) and abundance (b) and MGS richness (c) in fecal samples from 21+/-7-day-old neonates according to diet. (b-c) Data are displayed as box-and-whisker plots representing the interquartile range, with the central line indicating the median; individual data points are shown as dots. Statistical analyses were performed using the Fisher's exact test (a) and the Kruskal Wallis followed by the Wilcoxon rank sum test (b,c). ns: not significant; * p<0.05; *** p<0.001; **** p<0.0001. BF: breastfeeding; FF: formula feeding; MF: mixed feeding; MGS: metagenomic species.

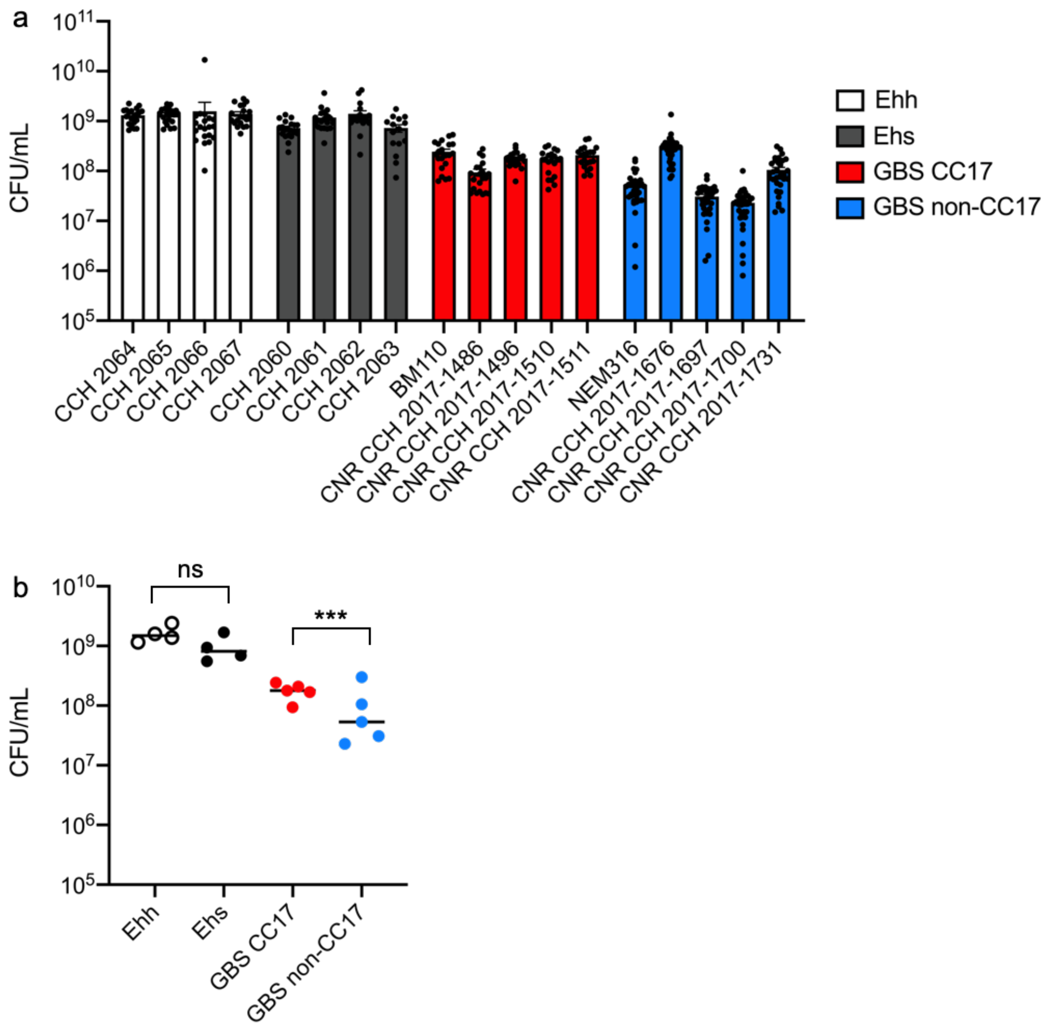


**Supplemental Figure 4. *E. hormaechei* and Group B *Streptococcus* (GBS) growth in co-culture in broth medium.** (a-b) Bacterial growth following 16h of co-culture showing the final colony forming unit (CFU) counts (a) and the mean final CFU counts (b) for each strain. Co-culture experiments were performed with *E. hormaechei* subsp. *hoffmannii* (Ehh) or *E. hormaechei* subsp. *steigerwaltii* (Ehs) in co-culture with GBS CC17 or GBS non-CC17. (a) Data are displayed as mean +/-SD and individual values for each experiment are represented as dots. (b) Data are displayed as mean CFU counts for each individual strain with line at median value. (a) Statistical analysis comparing CFU counts between strains of the same subgroup (Ehh, Ehs, GBS CC17, and GBS non-CC17) was performed using Kruskal-Wallis test and did not show any significant differences. (b) Statistical analysis comparing subgroups (Ehh vs. Ehs and GBS CC17 vs. GBS non-CC17) was performed using the Mann-Whitney test. Experiments were performed at least twice in duplicate. CC: clonal complex, Ehh: *E. hormaechei* subsp. *hoffmannii*; Ehs: *E. hormaechei* subsp. *steigerwaltii*; GBS: Group B *Streptococcus*. SD: standard deviation. ns: not significant; *** $p<0.001$.

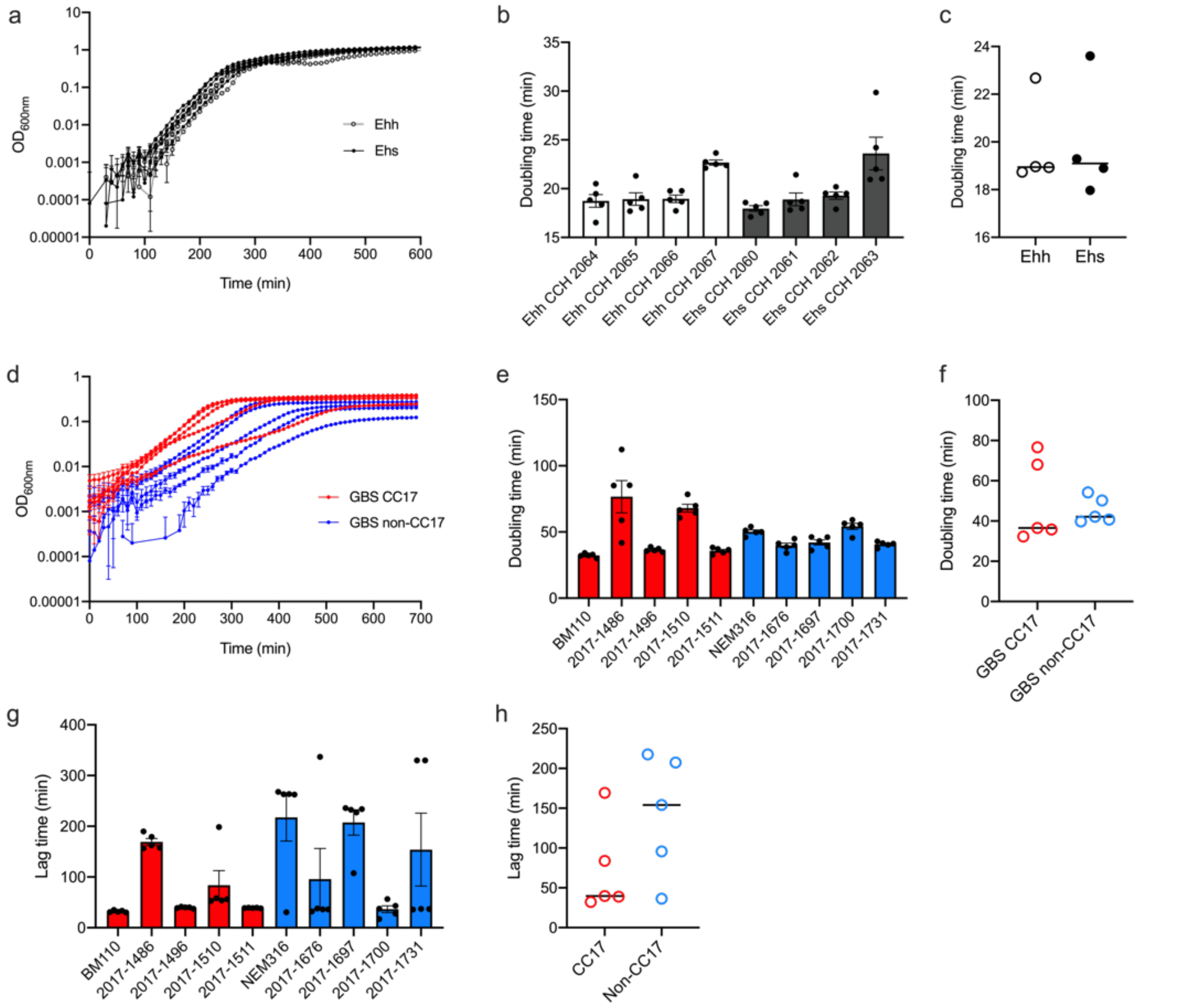


**Supplemental Figure 5.** ***E. hormaechei*** **and Group B Streptococcus growth characteristics in broth medium.** (a-c) *E. hormaechei* isolates growth curves (a), individual doubling time (b) and mean doubling time (c). (d-h) GBS isolates growth curves (d), individual doubling time (e), mean doubling time (f), individual lag time (g), and mean lag time (h). (b,e,g) Data are displayed as mean +/-SD; individual values for each experiment are represented as dots. (c,f,h) Data are displayed as mean value for each isolate with line at median. (b,e,g) Statistical analysis comparing strains of the same subgroup (Ehh, Ehs, GBS CC17, and GBS non-CC17) were performed using Kruskal-Wallis test and did not show any significant differences. (c,f,h) Statistical analysis comparing subgroups (Ehh vs. Ehs and GBS CC17 vs. GBS non-CC17) were performed using the Mann-Whitney test and did not show any significant differences. Experiments were performed at least twice in duplicate. CC: clonal complex, Ehh: *E. hormaechei* subsp. *hoffmannii*; Ehs: *E. hormaechei* subsp. *steigerwaltii*; GBS: Group B *Streptococcus*. SD: standard deviation.

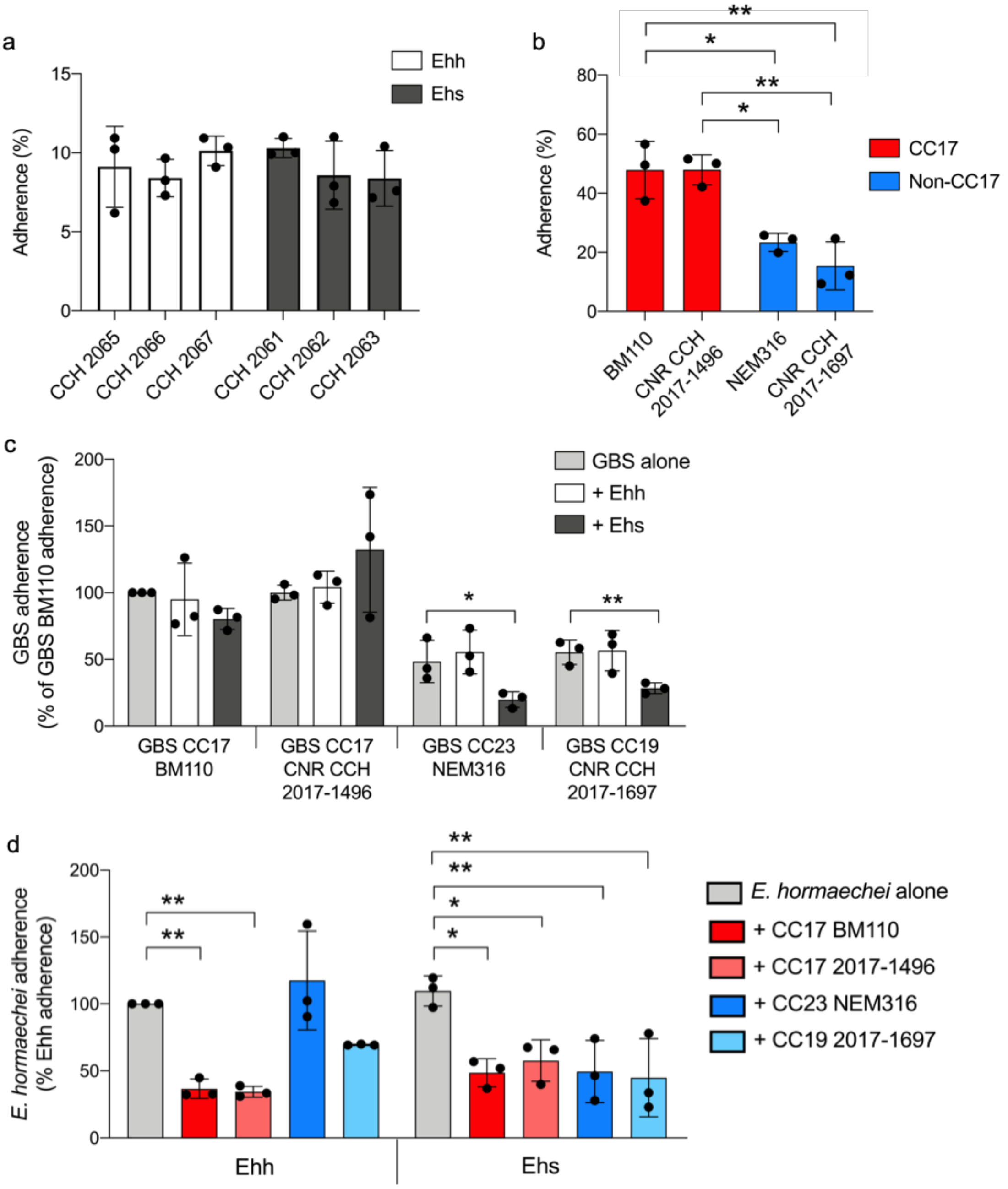


**Supplemental Figure 6. Adherence of *E. hormaechei* and Group B *Streptococcus* on Caco-2 enterocytes.** Cells were infected at a multiplicity of infection of 10 bacteria per cell. Following 2h of incubation, cells were washed and cell lysates were recovered for the quantification of adherent bacteria. Adherence of *E. hormaechei* and GBS isolates tested individually (a-b) and in competition experiments (c-d). *E. hormaechei* isolates used in competition experiments were Ehh CCH 2067 and Ehs CCH 2063 in a ratio GBS:*E. hormaechei* of 1:1. (c, d) Results are normalized to the adherence of GBS BM110 (c) and Ehh (d) alone. Experiments were performed three times in triplicate. Results are expressed as mean +/- SD. Dots represent the mean of triplicate values from individual experiments. Statistical analyses were performed using two-way ANOVA (a, b) and one-way ANOVA followed by Bonferroni post-test using GBS (c) or *E. hormaechei* (d) alone as control. * $p<0.05$; ** $p<0.01$. Ehh: *E. hormaechei* subsp. *hoffmannii*; Ehs: *E. hormaechei* subsp. *steigerwaltii*.
